# Formation of homolog pairing-induced domains in early *Drosophila* embryo genome

**DOI:** 10.64898/2026.02.04.703931

**Authors:** Lei Liu, Yanxia Jin, Yingyu Tao, Changbong Hyeon

## Abstract

Somatic homolog pairing is a defining feature of the diploid genome organization in *Drosophila* and underlies its transvection-based gene regulation. Here, to understand the physical effect of homolog pairing on the resulting three-dimensional (3D) organization, we employ the heterogeneous loop model and reconstruct 3D structures of the *Drosophila* embryo genome based on its haplotype-resolved Hi-C data. The resulting structures reveal robust end-to-end juxtaposition between homologous chromosomes amid substantial cell-to-cell variability. On sub-megabase scales, tight pairing between homologous loci at domain boundaries give rise to significant coincidence between *cis* and *trans*-homolog domain boundaries in the Hi-C map, while interior regions remain loosely associated. To uncover the physical origin of this organization, we compare the contact maps resulting from the polymer models implementing specific and non-specific button-mediated pairing mechanisms with Hi-C, finding that the intra-chromosomal contacts constrained by specifically paired inter-chromosomal buttons give rise to pairing-induced domains (PIDs). Our study suggests specific adhesive interactions as a central organizing principle of the diploid genome in *Drosophila* embryos.

**Significance:** Somatic homolog pairing distinguishes *Drosophila* from most eukaryotes; yet how the homolog pairing organizes *Drosophila* genome has remained elusive due to the lack of explicit model. By analyzing 3D structures reconstructed from haplotype-resolved Hi-C data, we clarify that specific homolog-recognizing buttons should generate pairing-induced domains that simultaneously organize *cis* and *trans*-homolog contacts. Our study provides a physical explanation on how a single molecular mechanism can simultaneously coordinate homolog pairing and architecture of chromatin domain in the diploid genome.

## INTRODUCTION

Most of eukaryotes are diploid. Their somatic cells carry two copies of genome inherited from each parent. Individual chromosomes, including homologous chromosomes, are segregated from each other, forming independent chromosome territories in the interphase, with their structure stabilized by intra-chromosomal contacts^1–3^. Single-cell Hi-C map and FISH images of human cells clarify that homologs in the interphase are segregated from each other and occupy their own territories inside cell nuclei^4,5^. In fact, for human, the coupling of two homologs could be highly deleterious, which necessitates antipairing mechanisms to suppress the homolog pairing in somatic cells^6^.

On the other hand, in some organisms, interchromosomal interactions bring homologs together, establishing somatic homolog pairings, which have to be discerned from those in meiosis characterized by strand invasion and recombination^7–9^ and from those during DNA break repair in the absence of an intact sister^10,11^. It has been known for *Drosophila*^6,12–15^ and yeast^16^ that robust homolog pairings play key roles in enabling the cross-homolog enhancer-promoter interaction termed transvection for gene regulation.

The haplotype-resolved Hi-C maps for somatic *Drosophila* cells^14,17^, which have recently been constructed through “phasing” of single nucleotide variants (SNVs), highlight significant *trans*-homolog (*thom*) contacts, suggesting the tight homolog pairing being achieved without incorporating DNA break as in the meiosis. However, as is common in many studies based solely on ensemble Hi-C^18^, the details of 3D genome architecture with homolog pairing are left unknown. The prominent features of *thom* domains observed in Hi-C map^14,17^and coincidence identified between *cis* (intra-chromosomal) and *thom* domain boundaries still await a clear physical explanation.

Several studies on somatic homolog pairing in *Drosophila* embryo genome have hypothesized a mechanism that interactions between homologous “buttons” interspersed along the chromatin fiber create the end-to-end juxtaposition of two chromosomes^14,17,19–23^. To extract the density of buttons along the chromatin and the strength of the specific inter-button interaction, a polymer-based simulation was performed and its results were compared with the live imaging of chromosomal loci demonstrating homolog pairing^20^. In another study^21^, it was argued that, even if the individual button interactions are non-specific, two homologs with non-uniformly distributed barcode-like buttons can still achieve end-to-end alignment since such a configuration is energetically more favorable than a random alignment with many misatches.

While these theoretical studies focus on how likely or how fast homologs are paired^8^, neither of them have addressed its impact on shaping the real 3D organization of chromosomes. While the button mechanism at present is largely phenomenological, it is at least known that the button sequences are enriched around *cis* domain (or topologically associated domain, TAD) boundaries^14^ that accommodate various architectural proteins (APs, a.k.a insulator proteins, such as CTCF, BEAF-32, CP190 and Su(Hw)), each of which binds a unique DNA motif^14,20,22,24,25^.

Here, in order to have a concrete understanding of the higher-order genome organization associated with somatic homolog pairing, we apply the heterogeneous loop model (HLM) approach that some of us developed^26–30^ to haplotype-resolved Hi-C maps and reconstruct 3D structures of *Drosophila* embryo genome. By comparing the Hi-C-inferred chromosome structures exhibiting somatic homolog pairing with those predicted by the diametrically different two button models described above, we aim to gain new insights into the *Drosophila* genome architecture induced by somatic homolog pairing.

## MATERIALS AND METHODS

### Hi-C, TADs and ChIP-seq data

Hi-C and Micro-C datasets were collected from the GEO repository. The contact frequencies between loci pairs were counted at a prescribed resolution, followed by normalization using the Knight-Ruiz method^31^ with Juicer^32^, Cooler^33^, or in-house codes, depending on the format of the data. Next, dangling segments were eliminated from our analysis if *>*90% of their intrachromosomal contact frequencies were zero.

First, for intra-chromosomal pairs, the arc length (*s*)-dependent mean contact probability is defined as

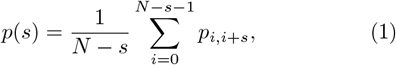

with *N* the number of segments on that chromosome. Then, one can quantify the extent of enrichment or depletion of contacts between (*i, j*) loci pair by rescaling the contact probability *p*_*ij*_ with the mean intra-chain contact probability *p*(*s*) when the two segments are in the same chain and with the mean inter-chain contact probability 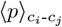 where *c*_*i*_ denotes the chromosome (arm) to which the *i*-th segment belongs,

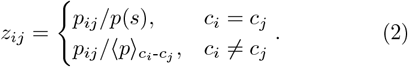

For the genomic regions corresponding to TADs, we directly used genomic coordinates of TADs defined in early embryos^34^, in third instar larval eye discs^19^, and in S2R+ cells^35^ derived from late embryos in *Drosophila*, which specify 2023, 1122, and 4123 TADs, respectively. The reMap2022 database^36^ was used for analyzing the correlation between protein binding and homolog pairing. It standardized and integrated 1205 public ChIP-seq datasets across 550 DNA-binding proteins in *Drosophila*, among which 183 datasets were selected from embryos. While the Hi-C reads were mapped to the *Drosophila* reference genome dm3, the Micro-C and reMap2022 datasets were built based on a more recent *Drosophila* genome assembly dm6. We chose dm3 as a reference in this study, and converted data between dm3 and dm6 by using the UCSC Genome Browser utility liftOver^37^ when necessary.

### Construction of 3D structural ensemble of fly genome

3D structural ensembles are constructed based on Hi-C maps by using HLM, which models the whole genome as *M* polymer chains composed of total *N* monomers (sites), each representing a chromatin segment of a prescribed genomic length^30^. The energy potential of the model is defined as the sum of harmonic restraints between all pairs of genomic segments, that effectively maps the 3D structure of a chromatin fiber or the whole genome onto *M* vulcanized Gaussian polymer(s)^38–40^. The strength of each harmonic restraint is iteratively adjusted until the contact probabilities 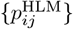 converges to those of Hi-C map, 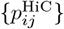 The model enables one to clarify the structural origin of the variable gene expression level^26^, to visualize structural changes along the cell cycle, to predict multi-way chromatin contacts^28^, and to address the nature of various experimental methods for mapping 3D genome architecture2^9^. Thus, it adds a new dimension to the current genome research that relies solely on Hi-C without explicit structures^2,41-43^.

Specifically, the effective energy potential for the model is written as a sum of harmonic restraints between the genomic segments,

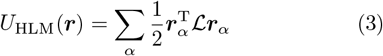

where ***r***__*α*__ = (*r*_0, *α*_, *r*_1, *α*_, *r*_2, *α*_, …, *r*_*N−*1, *α*_) with *α* ∈ (*x, y, z*), and the summation over (*i, j*) includes both intra- and inter-chain monomer pairs. The Laplacian matrix ℒ is defined asℒ= 𝒟− 𝒦, where is a stiffness matrix for the strength of harmonic restraint between *i* and *j* segments *k*_*ij*_ = (𝒦)_*ij*_ (*i* ≠*j*) and 𝒟 is a diagonal matrix of elements 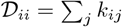^28,29^.

By considering the eigenvalue decomposition the matrix ℒ, the covariance matrix Σ can be calculated as

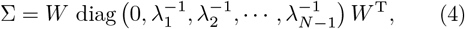

where the *i*th column vector of the matrix *W* and *λ*_*i*_ are the corresponding *i*th eigenvector and eigenvalue of the matrix ℒ, respectively. For the given Σ, it is straightforward to calculate the pairwise contact probability:

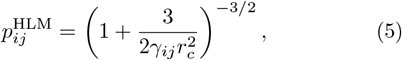

where *r*_*c*_ corresponds to the effective capture radius of the cross-linking agent, and

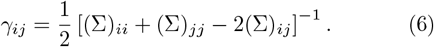

A procedure of constrained optimization, which minimizes the difference between 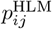 and 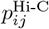, offers the optimal parameters for the model 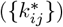^28^.

Once 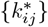 are decided, a genome structure can be generated in two steps^29,44^. First, we calculate the normal coordinates, *X*_*i,α*_, by using

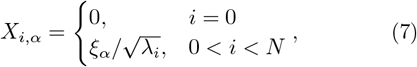

where the random variable *ξ*_*α*_ obeys the normal distribution with ⟨*ξ*_*α*_⟩ = 0 and ⟨*ξ*_*α*_*ξ*_*β*_⟩ = *δ*_α*β*_. Next, the normal coordinates are converted to the cartesian coordinates of chromatin segments by

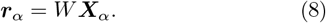

Repeating this procedure ∼10^4^ times, we generated an ensemble of 3D genome structures.

### Button model for homolog pairing

We adapted the polymer model developed by Marshall and Fung^21^, so as to assess chromosome structures on a short genomic scale (*<* 1 Mb) resulting from the polymer model with specific buttons^20^ and those from another model with non-specific buttons^21^. We modeled two pairs of chromatin fibers using worm-like chain^45^, with each fiber consisting of 90 monomers representing a genomic region of 900 kb, and specified the genomic positions of pairing buttons along the fiber. First, the distance between adjacent monomers (*r*) is constrained by a harmonic potential

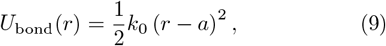

with a length unit of *a* and a spring constant of *k*_0_ = 25 *k*_*B*_*T/a*^2^. Second, the chain flexibility is modulated by

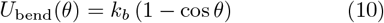

where *k*_*b*_ is the bending parameter and *θ* is the angle between consecutive bonds along the chain. To mimic the indirect effect of centromere clustering on pairing, we restrain the coordinate of the first monomers of four chains 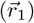 around a center of the coordinate system using a restoring potential,

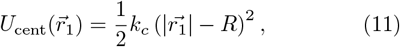

with *k*_*c*_ = 35 *k*_*B*_*T/a*^2^ and *R* = 2*a*.

Two-stage Brownian dynamics simulations were performed to sample the conformations of homologs. During the first stage, four polymer chains were thermalized from random coils by integrating the following equation of motion,

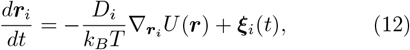

where *U* (***r***) equals the summation of the three terms above (Eqs. 9-11), *D*_*i*_ is the bare diffusion coefficient of the *i*-th monomer, and ***ξ***_*i*_(*t*) denotes the Gaussian noise satisfying ***ξ***_*i*_(*t*) = 0 and ***ξ***_*i*_(*t*) ***ξ***_*j*_(*t*^*I*^) = 6*D*_*i*_*δ*_*ij*_*δ*(*t −t*^*I*^).

Next, button-based mechanisms were switched on to model somatic homolog pairing. Each integration step consists of three substeps: (i) Currently paired buttons were unpaired with a chance of *p*_*u*_ (= 0.001). (ii) The coordinate of each monomer in the four chains was integrated and updated according to Eq. 12, whereas any two paired button monomers were restrained to move together. (iii) Any two unpaired buttons in a spatial proximity (*r < r*_*p*_ = *a/*2) are considered to form the paired state. The non-specific button model allows a button to pair with any other buttons (on the same or different chains)^21^, whereas the specific button model only allows pairing between homologous buttons^20^.

Simulations in the pairing stage were performed for 8.1*×*10^5^ *τ*_BD_ with an integration time step of *δt* =0.01 *τ*_BD_, where *τ*_BD_ = *a*^2^*/D* is the Brownian time of monomers. Conformational properties were calculated based on structures from the second half of this stage, when the fraction of *correctly* paired buttons (*f*_pair_) became stable. The chain stiffness parameter *k*_*b*_ in Eq. 10 was varied from 0 and 9 in the nonspecific model, and was set to 0 in the specific one. For each setting, 64 independent runs were generated. Although it has been shown that the parameters, *R, p*_*u*_, and *D*_*i*_, affect the pairing success and dynamics^21^, we focus on the structural properties of paired homologs while fixing those parameters to specific values.

## RESULTS

Genome architecture in *Drosophila* has been extensively studied^47–49^. In early embryos, their cell nuclei are diploid, consisting of four chromosome pairs, three autosomes (chr2, chr3, and chr4) and one sex chromosome (chrX). On the other hand, tetraploid cells accumulate in the adult *Drosophila* brain, which is thought to protect against the aging related cell damage^50^. The tight association of the somatic homolog pairing is the most dramatic feature in the *Drosophila* genome organization that begins to develop during the interphase cycle 14 in embryogenesis^51,52^. Furthermore, recent studies^53,54^ indicate that, in early embryos, topologically associated domains (TADs) are organized through architectural protein (AP) mediated-homolog pairing, rather than loop extrusion mechanism that dominate genome organization in many other species^55^. In what follows, we will rationalize a coupling between *trans*-homolog interactions and *cis*-domain formation by analyzing 3D structural models of *Drosophila* genome reconstructed from HLM.

### 3D architecture of *Drosophila* genome

We employ the non-haplotype-resolved genome-wide Hi-C data from Kc_167_ cell line^46^ as an input to the HLM approach, and analyze the resulting architecture of *Drosophila* genome acquired to extract the overall structural properties. Notably, the contact map (Fig. 1A), the arc length (*s*)-dependent mean intra-chromosomal contact probability (Fig. 1B), and the pairwise contact probability (Fig. 1C) obtained from the model are in excellent agreement with those directly calculated using the Hi-C map. The HLM-based 3D structures indicate that two metacentric autosomes, chr2 and chr3, are folded nearly in half with respect to their centromeres, giving rise to the left (2L, 3L) and the right (2R, 3R) arms, and this feature is reflected as the anti-diagonal bands in the contact maps (Fig. 1A).

**FIG. 1.**
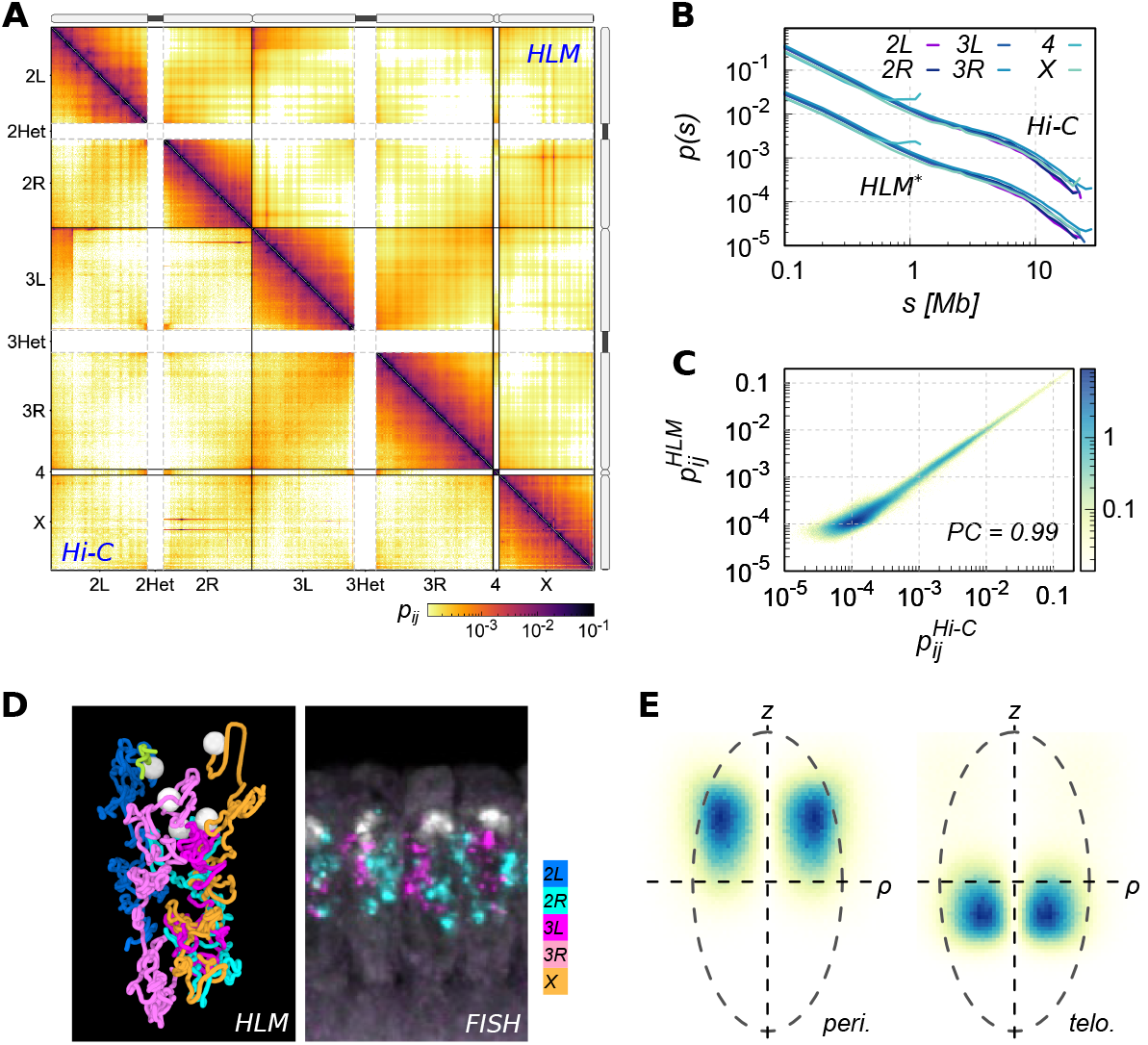
3D architecture of *Drosophila* genome. (**A**) Contact probabilities of *Drosophila*_167_ cells from Hi-C (upper triangle) and HLM (lower triangle)^46^. (**B**) Mean intra-arm contact probability, *p*(*s*), for 2L, 2R, 3L, 3R, 4, and X. The result for HLM is shifted vertically (divided by 10) for the sake of visual clarity. (**C**) Genome-wide correlation between Hi-C and HLM. (**D**) A representative model structure from HLM (left), compared to a 3D-DNA FISH image of *Drosophila* embryos taken from Ref. (47) where only chr2R and chr3L are stained (right). Different colors label different chromosomes, and the white spheres in the model represent pericentromic chromatin segments. (**E**) Distributions of pericentromeric and telomeric regions determined from the structural models generated by HLM.

Around the principal axis defined using the pericentromeres and telomeres^48^, the predicted 3D model is characterized by the prolate shape with an aspect ratio of 2 (Fig. 1D and E). The four chromosome arms (2L, 2R, 3L, and 3R) and chrX adopt extended conformations along the principal axis of the enveloping ellipsoid, and the pericentromeric regions (white spheres in Fig. 1D) and telomeric regions are clustered together in the opposite hemispheres of the nucleus (Fig. 1E). Previously, depletion of heterochromatin protein 1a (HP1a) was shown to disrupt the centromere clustering^46^, leading to loss of the so-called Rabl configuration^56^. Both centromere and telomere clusterings are known to facilitate homolog pairing^21,57, 58^ .

A few qualitative differences are found between the structure predicted from our approach (Figs. 1D, E) and those predicted in Ref. (59). The positions of pericentromeres are more dispersed, and telomeres are closer to the nuclear center in HLM. We, however, note that these differences may arise from the artificial restraints imposed on the chromosomes in Ref. (59), specifically those among the pericentromeres and between telomeres and nuclear envelope. A spherical nuclear shape is also presumed in their modeling. Furthermore, instead of using the whole genome Hi-C map, chromosomes were built one by one based on the Hi-C map of each chain. The whole genome structure in Ref. (59), obtained by assembling the individual chromosome chains under the aforementioned restraints, is at odds with our 3D model that better aligns with the FISH image (Fig. 1D, right panel).

### Genome-wide juxtaposition of homologs

Haplotype-resolved Hi-C maps are obtained from Hi-C sequencing data processed through the phasing of heterozygous variants. The Hi-C reads are aligned to the reference genome, and each read end is assigned to the maternal or paternal haplotype based on its overlap with phased SNVs. Contacts where both ends are assigned to the same haplotype correspond to *cis* maternal or *cis* paternal maps, while *trans*-homolog (*thom*) or *trans*-heterolog contact maps are obtained when the two ends are assigned to different haplotypes.

The extensive *trans*-homolog interaction is a hallmark of *Drosophila* genome^60^. Homolog pairings and allelic-specific genome structures are associated with many cellular processes^6^. According to the haplotype-resolved Hi-C^17^, strong similarities are observed between the contact maps of the maternal and paternal genomes and those of trans-homolog interactions in early *Drosophila* embryos (compare the Hi-C map regions enclosed in the red and blue squares in Fig. 2A). This pronounced similarity is contrasted against the *trans*-heterolog contacts, which exhibit substantially weaker interaction intensities (Fig. 2A).

**FIG. 2.**
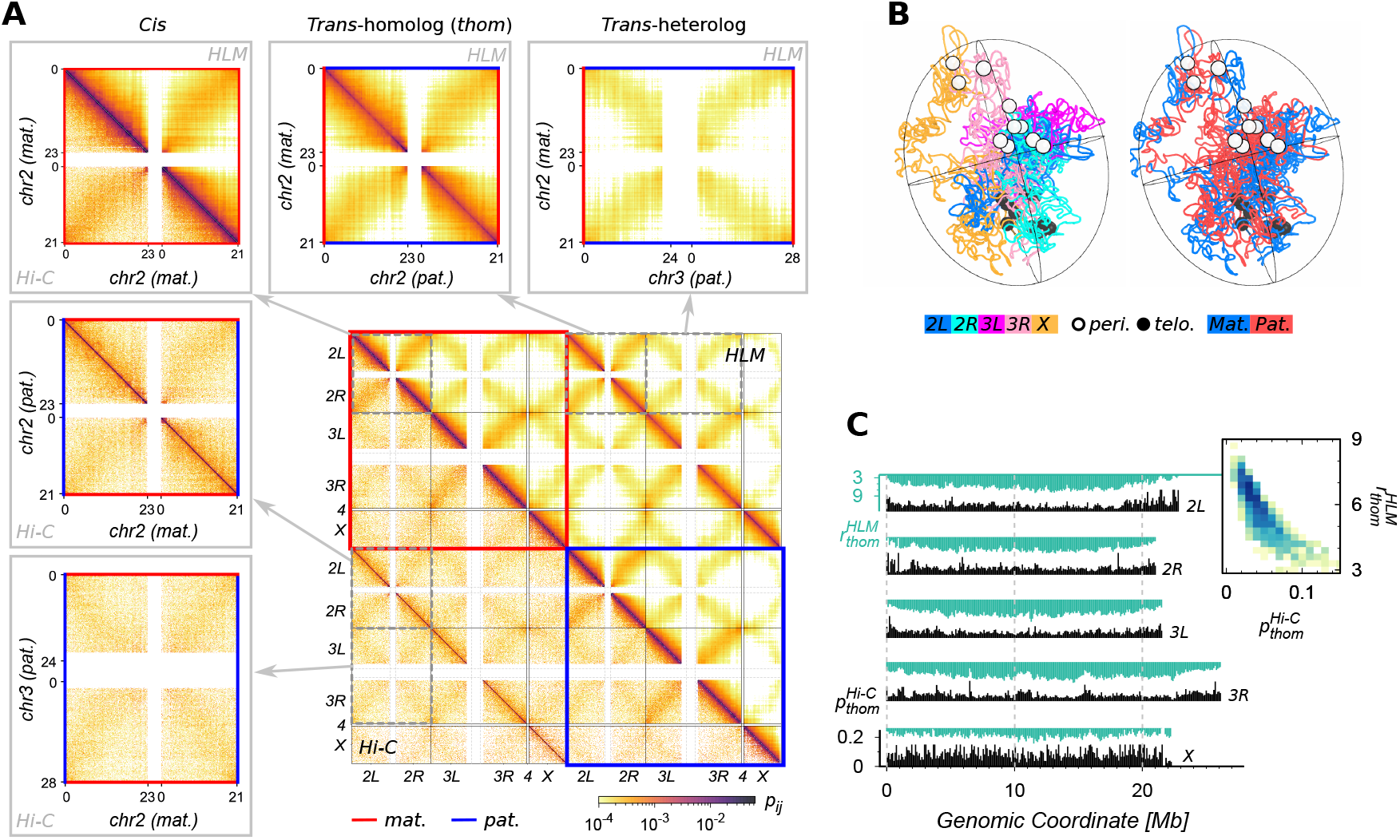
Modeling diploid genome of early *Drosophila* embryos. (**A**) Genome-wide contact probability matrix from HLM (upper triangle) versus that from haplotype-resolved Hi-C^17^ (lower triangle). In the main contact map, the contacts originating from the maternal and paternal genomes are demarcated by red and blue boxes, respectively. Zoomed-in maps of intra-chromosomal contacts of maternal chr2 (*cis* Mat), inter-chromosomal contacts between maternal chr2 and paternal chr2 (*trans*-homolog), and those between maternal chr2 and paternal chr3 (*trans*-heterolog) are enclosed in gray boxes. (**B**) A representative HLM-generated 3D genome structure that depicts the 5 chromosome arms (left) and maternal/paternal chromosome (right). (**C**) Profiles of *trans*-homolog contact probability from Hi-C 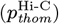 and the mean distance from HLM 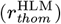 binned at 100 kb,The inset shows the scatter plot of 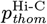 versus 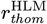.

Our 3D genome structures reconstructed at 100 kb resolution allow us to directly evaluate the tightness of homolog pairing, which is lacking in studies relying entirely on Hi-C^14,17^. Again, the agreement of the mean intra-arm contact probabilities (*p*(*s*), Fig. S1A), contact probabilities (*p*_*ij*_, Fig. S1B) as well as the contact map (Fig. 2A) derived from HLM with those from Hi-C is remarkable. The extent of pairing between homologous chromosomes is visualized in a representative HLM-generated 3D model (Fig. 2C). Despite great cell-to-cell variations in the 3D genome (see Fig. S1C), the overall similarity between the maternal and paternal genome organizations is persistent within *each* cell.

The profiles of *trans*-homolog contact probability, *p*_*thom*_, provide further details of how tightly homologous segements are paired in each chromosome (Fig. 2C). Except for the relatively larger values of *p*_*thom*_ at both ends of chromatin chains involving the clusterings of centromeres and telomeres, the extent of homolog pairing along the chain is non-uniform. Nevertheless, the mean distances between allelic segments extracted from HLM 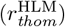 exhibit clear anti-correlation against 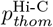 with a genome-wide Spearman correlation of 0.86 (see inset of Fig. 2C).

### Organization of homologous chromosomes at the TAD scale

Hi-C maps acquired at higher resolution further clarify the variations of pairing between two homologs. In Fig. 3A, the lower triangle demonstrates the haplotype-resolved Hi-C contact probability matrix for a 900 kb genomic region on chr2R from PnM cells of *Drosophila* embryos^14^ at 10 kb resolution. This dataset reveals ∼7.8 times more number of *trans*-homolog contacts ^14^ (see also Fig. S2A) than that of typical Hi-C map for *Drosophila* embryo cells.

**FIG. 3.**
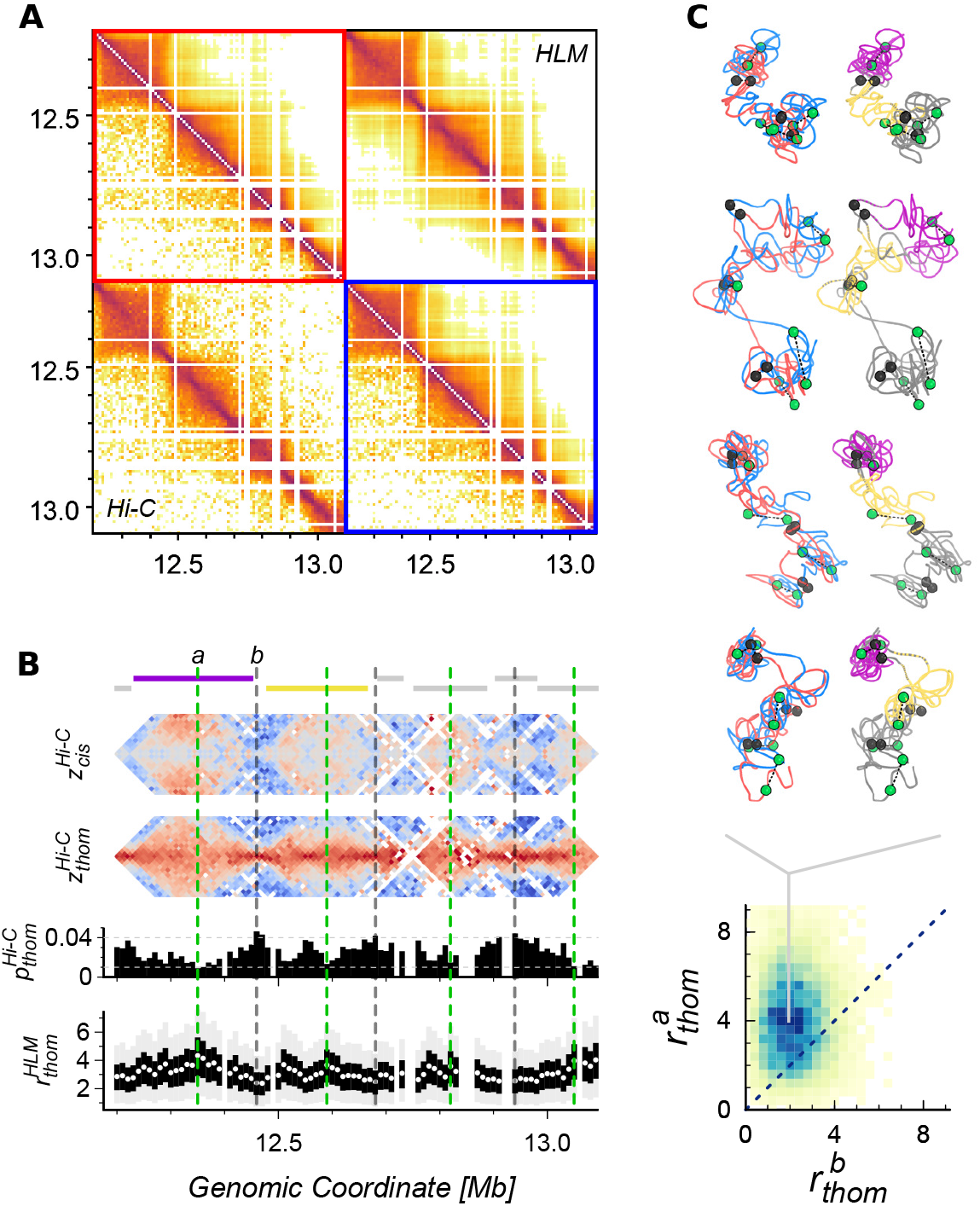
Homolog structure on a sub-megabase scale. (**A**) contact map from HLM (upper triangle) versus that from haplotyperesolved Hi-C^14^ (lower triangle) for the genomic region of chr2R:12,200,000-13,100,000. (**B**) Profiles of *cis* TADs^34^, *cis* and *trans*-homolog contact enrichment 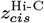 and 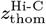, *trans*-homolog contact probability 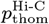and HLM-based *trans*-homolog distance 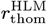 . The genomic regions of TADs are demarcated with the horizontal bars on the top. (**C**) Distributions of the *trans*-homolog distance at the genomic loci *a* and *b* in HLM-generated structural ensemble. Shown on the top are the representative structures depicted in two different ways: (left) maternal and paternal chromatins in red and blue, (right) the two largest TADs in purple and yellow. The four green and three black pairs of spheres depict the loosely and tightly paired loci, respectively, whose genomic positions are marked in (**B**) with vertical dashed lines.

Of particular note are the domain structures in the *trans*-homolog (*thom*) contacts and the coincidence of *cis* and *thom* domain boundaries. These properties can be better discerned using the parameter (*z*_*ij*_) defined in Eq. ^2^ that measures the enrichment or depletion of contacts (Fig. 3B). Based on the *cis* TADs called from *Drosophila* embryos^34^ (top bars in Fig. 3B), 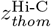 is dramatically reduced inside the domains. The genomic loci corresponding to the domain boundaries, whose positions is marked using black vertical dashed lines, are characterized with the three highest values of 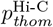and the smallest value of 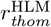 indicating that homolog pairings in the domain boundaries are tighter than other parts. This is consistent with the genome-scale analysis on pairing of loci with respect to domain boundaries (Fig. 2g in Ref. (14)), as well as our statistical analysis of *trans*-homolog contacts based on the pairing score (Fig. S2C).

The *trans*-homolog distance based on HLM-generated 3D structures,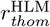, are almost perfectly anti-correlated with 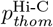 (Fig. 3B, Spearman correlation−0.99). We analyzed the structural ensemble giving rise to TAD domains based on the value of 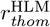. For example, at the genomic locations of *a* and *b* labeled at the top of Fig. 3B, the former and the latter denoting the center and the boundary of the largest TAD, respectively, are characterized with the lowest and the highest value of 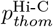. As shown in the two-dimensional histogram of inter-homolog distances at *a* and *b* together with some of the representative chromatin structures (Fig. 3C), the domain center (*a*) has a greater inter-homolog distance than the domain boundary (*b*) in majority (85 %) of the HLM-based structures.

### Contact maps from non-specific button model

As visualized with HLM-based structures, the effect of tightly paired loci (buttons) on chromatin architecture is two fold: (i) association of homologs and (ii) organization of *cis* and *thom* domains. To understand the interplay between the two processes, we study both the specific and non-specific button mechanisms (Fig. 5A) by performing Brownian dynamics simulations of the homolog pairing (Materials and Methods). Specifically, we consider a system consisting of two pairs of (semi)flexible polymer chains. each of which represents a genomic region of 900 kb, with 90 monomers. Two button sequences with a line density of 0.3 are selected based on the Micro-C contact maps of chr2/3R:12,200,000-13,100,000 (Fig. 5B), and then they are assigned to two pairs of chains *m*_1_ and *p*_1_, *m*_2_ and *p*_2_, respectively. For each sequence, the largest gap (consecutive monomers without buttons) and trail (a genomic region with high button density) are marked with a grey and black bar, respectively, in Fig. 5B.

For non-specific button model, we tuned the chain flexibility, which has previously been shown to be a key determinant of the success of homolog pairing^21^. By increasing the parameter for bending rigidity *k*_*b*_ (Eq. 10), the persistence length *l*_*p*_ increases from 0 to 8*a* (Fig. S3A). As shown in Fig. 4C (see also Supporting Movies 1-3), at small *l*_*p*_ (*<* 4*a*), the two pairs of homologs effectively behave like collapsed random coils while forming buttons at random positions, which results in a negligible fraction of correct pairings (*f*_pair_≈ 0). The correct buttons (or homolog pairings) are made with high fidelity only if rigidity of chains is greater than a certain threshold (*l*_*p*_≈ 5.5*a*).

**FIG. 4.**
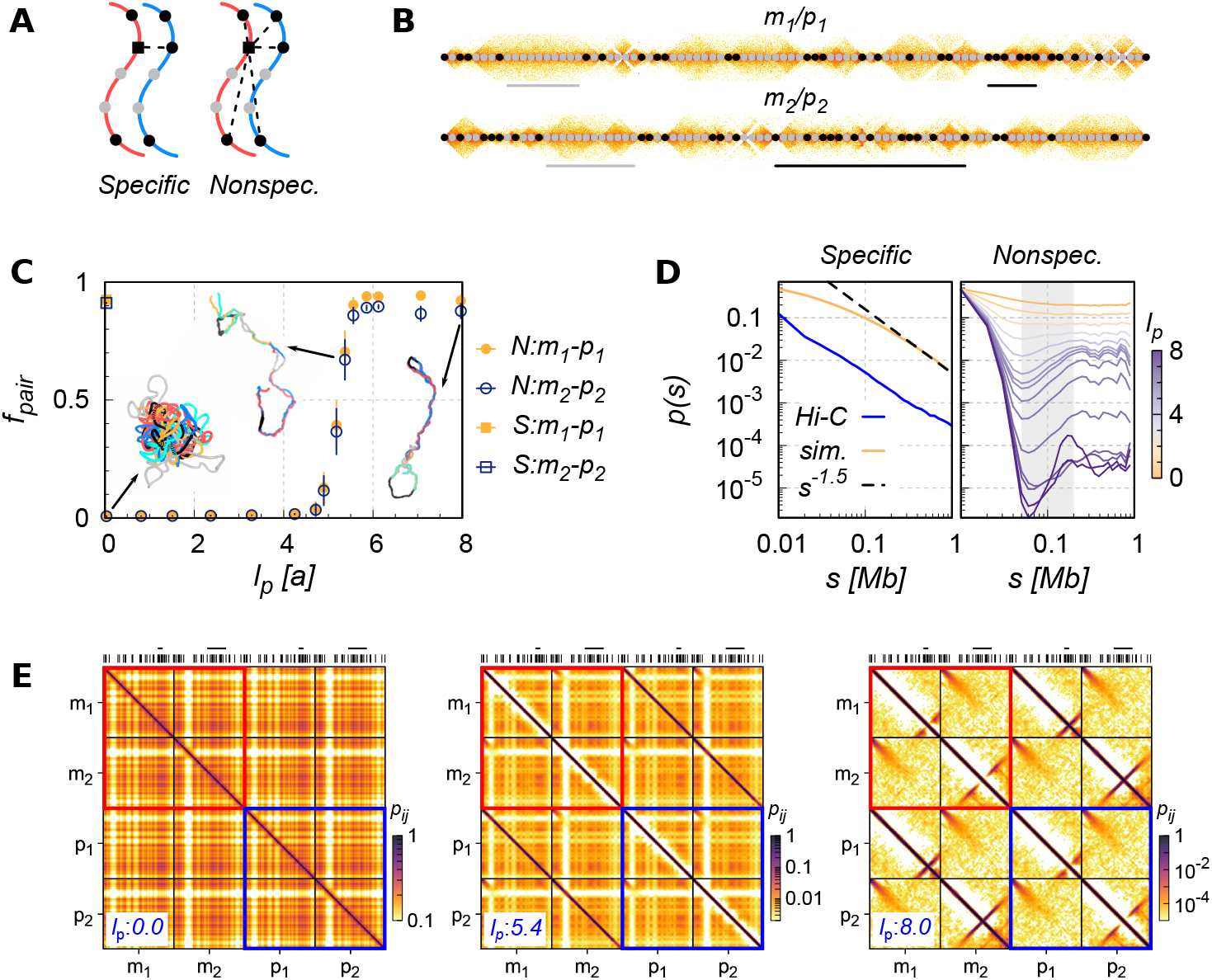
Contact maps from nonspecific button model. (**A**) An illustration of specific versus non-specific button mechanism. A squared button can only pair with its homologous button in the former, while it is allowed to pair with any other button (on the same chain or a different chain) in the latter. (**B**) The buttons (black dots) on chains *m*_1_*/p*_1_and *m*_2_*/p*_2_, which were chosen based on Micro-C data of two 900 kb regions on chr2R (top) and chr3R (bottom)^34^, respectively. The gray and black bars label the largest gap and trail for each button sequence. (**C**) The fraction of correctly paired buttons (*f*_pair_) as a function of the persistence length of single chain (*l*_*p*_), in specific (S) or non-specific (N) models. Three representative conformations at *l*_*p*_ = 0.0, 5.4, and 8.0 *a* are shown as insets, which are not plotted using the same scale (see Fig. S3B). Chains *m*_1_, *p*_1_, *m*_2_, and *p*_2_ are colored in red, blue, cyan, and orange, respectively. (**D**) Mean intra-chain contact probability from haplotype-resolved Hi-C^14^ and from modeling. (**E**) Contact maps at three different values of *l*_*p*_ with the button barcode on top.

The mean intra-chain contact probability, *p*(*s*), calculated using the conformational ensemble of this model shows features at odds with Hi-C (Fig. 4D). The intra-chain contact probability *p*(*s*) decreases non-monotonically for *l*_*p*_ ≳2*a*. Such trend is amplified as *l*_*p*_ increases further, which apparently differs from the Hi-C-based *p*(*s*) which decreases monotonically on the scales of interest. The non-monotonic variation of *p*(*s*) at small *s* can be explained by the competition between the bending penalty and the entropic cost associated with the loop formation of a semiflexible chain with an increasing arc length. It has been shown for semi-flexible chain that *p*(*s*) is maximized when *s/l*_*p*_ ∼ (2− 3)^61–63^.

The contact maps calculated using the ensemble of assemblies made of the four chains *m*_1_, *m*_2_, *p*_1_, and *p*_2_ for different values of *l*_*p*_ offer further physical insight into the resulting chain organization (Fig. 4E). For flexible chains (*l*_*p*_ = 0), the contact map shows a plaided pattern in alignment with the button sequence (the leftmost panel of Fig. 4E). Uninterrupted horizontal and vertical stripes with small *p*_*ij*_ in the contact map are formed around the location of button gaps, indicating that the chain segments in the gaps rarely interact with others. As the chains get stiffer (e.g., *l*_*p*_ = 5.4*a*), a contact-depleted zone is also formed around the main diagonal of the map, and its width increases with *l*_*p*_ (the middle panel of Fig. 4E). This is consistent with the non-monotonic variations of *p*(*s*) at small *s* (Fig. 4D). For *l*_*p*_ = 8.0*a*, hairpin-like structures are formed around the regions replete with buttons, which contributes to formation of the short anti-diagonal stripes in the contact map (the rightmost panel of Fig. 4E)^64^. None of the above characteristics can be found in Fig. 3A. Thus, regardless of the extent of homolog pairing, their 3D chain organization in non-specific button model is not compatible with Hi-C map.

### Pairing-induced domains organized by specific buttons

In contrast to the foregoing section addressing the outcomes from non-specific button mechanism, the specific button mechanism can produce the main features observed in the haplotype-resolved Hi-C data of *Drosophila* (Fig. 4A). Homologous polymer chains are successfully paired for *f*_pair_ ≥0.9 (Fig. 4C) even when *l*_*p*_ = 0 (Supporting Movie 4). The contact probability *p*(*s*) (left panel in Fig. 4D), displays a power law decay of *p*(*s*) ∼*s*^−1.5^ at large values of *s*, which is also consistent with the analysis using Hi-C map.

*Trans*-homolog contacts (*p*_thom_) resembling the *cis* contacts are manifested in the contact map (see the *m*_1_-*p*_1_ or *m*_2_-*p*_2_ region of the contact map and compare it with the *cis*-domain corresponding to the region of *m*_1_-*m*_1_ or *m*_2_-*m*_2_ in Fig. 5A), the organization of which is reflected in a representative conformation of polymer chains displaying the end-to-end juxtaposition between two homologous chains (see the inset of Fig. 5A). As shown by *z*_thom_ that quantifies the enrichment of the *trans*-homolog contacts between *m*_1_ and *p*_1_, *p*_thom_ of each monomer depends on its position along the chain, which is negatively correlated with the arc distance to its nearest button (Fig. S4C). Therefore, *p*_thom_ and *z*_thom_ can be much lower in the interior than at the boundary of *thom* domains.

**FIG. 5.**
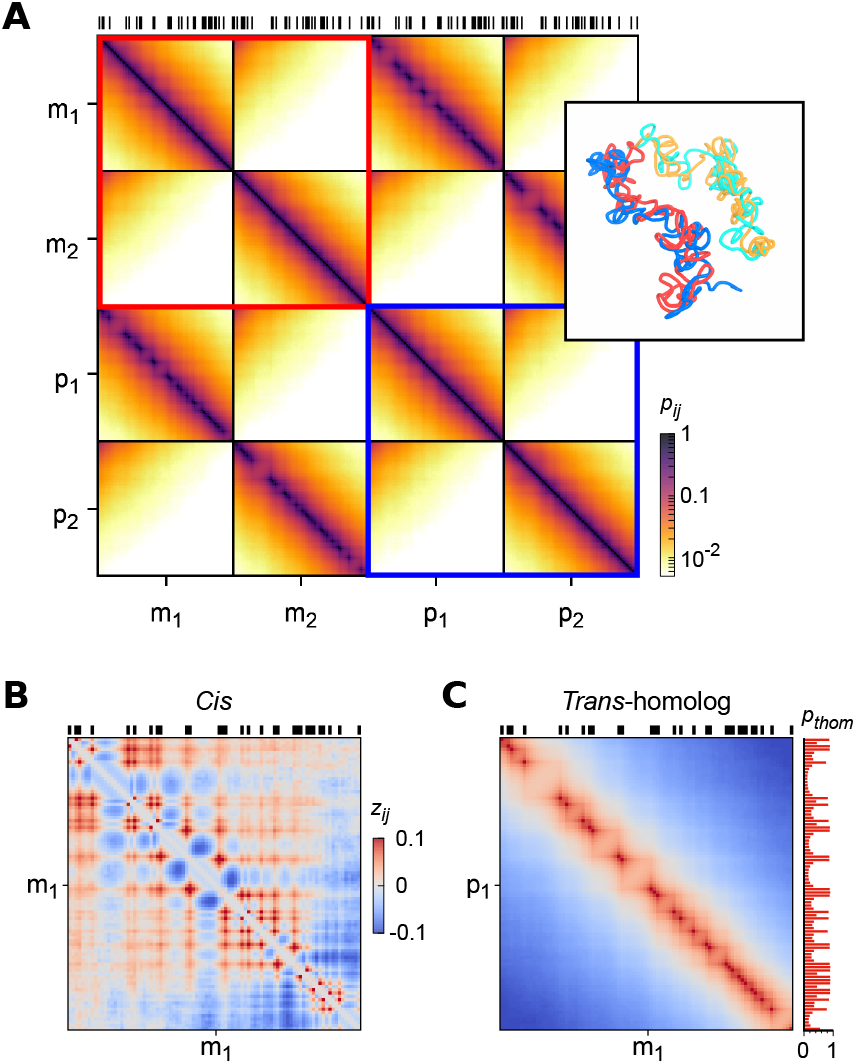
Contacts from specific button model. (**A**) Contact prabability map calculated using 3D conformations of the specific button model, with the button barcode on top and a representative structure in the inset. The enrichment maps of *z*_*ij*_ for (**B**) *cis* and (**C**) *trans*-homolog contact between chains *m*_1_ and *p*_1_ calculated using Eq. 2. Nearly identical results from the other pair of homologous chains (*m*_2_-*p*_2_) are demonstrated in Fig. S4.

## DISCUSSION

One of the most interesting findings from the specific button model is the formation of *cis* domains flanked by two buttons (Fig. 5B) whose boundaries display perfect overlap with the boundaries of *thom* domains. To better understand the *cis* domain formation in the specific button model, we consider a simplified case in which all homologous buttons are paired, resulting in a pairing-induced structure with many bubbles (Fig. 6A), each of which is made of two arcs with different parental origin. Now, we compare the contact probabilities between points (*i, j*) and (*m, n*) that are located on the same (blue) chromosome and are separated by the same genomic distance of *s*, i.e., |*i*−*j*|= |*m* −*n*| = *s*, with a setting that while *m* and *n* are in the same bubble, *i* and *j*, interrupted by an button point *o*, are separated into two neighboring bubbles. Under this setting, the intervening buttons are expected to insulate *cis* contacts, giving rise to asymmetric contact probabilities (*p*_*ij*_ *< p*_*mn*_).

**FIG. 6.**
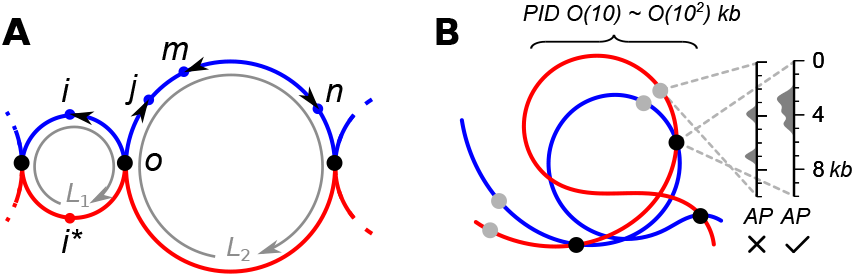
Pairing-induced domain structure. (**A**) An illustration of a bubble chain to account for the *cis* domain formation.(**B**) A proposed hierachical organization of somatic homologs in *Drosphila* on a sub-megabase scale. The differential distribution of APs determines the specificity of the adhesion between homologous loci. As opposed to the gray buttons characterized with sparse AP distribution, the adhesive, specific buttons (black dots) induce pairings and yield contact domains.

Our idea of the pairing-induced domain (PID) structure can be further elaborated by considering a bubble chain, where every homologous button pair is merged to a monomer, giving rise to a contact map nearly indistinguishable from that of the specific button model (Fig. S5). More quantitative discussion, employing an idea of polymer physics^65–67^, of how *cis* and *thom* contacts are modulated in the presence of an intervening button and the origin of overlap between *cis* and *thom* domain boundaries, is provided in the Supporting Material. Based on the generic nature of the contact-insulating function of buttons, pairing of sister chromatids should contribute to contact domains in replicated genome as well^68–70^, although some of the key molecular players in sister-chromatid pairing, e.g., cohesin, are dispensable for somatic homolog pairing^71–73^.

Analysis of the haplotype-resolved Hi-C data of *Drosophila* embryos shows that somatic homolog pairing in epigenetically active state is tighter (Fig. S2D and see also the results based on PnM cells found in Fig. 3b of Ref. (14)). Collecting all ChIP-seq datasets involving *Drosophila* embryos from reMap2022 database^36^, we calculated the correlation between ChIP-seq peak density and *trans*-homolog contact probability. According to Fig. S2B and Table S1, besides the pioneer transcription factor Zelda^17^, several architectural proteins (APs), such as upSET, CBP and Rsf1, may promote somatic homolog pairing. The small value of the highest correlation coefficient (*<* 0.3; see also Table S3 in Ref. (14)),however, calls for further studies to pin down the APs responsible for pairing.

As shown in Fig. 4, the non-specific button model does not produce a contact map compatible with Hi-C map on the physically relevant genomic scales from 0.01 to 1 Mb. We also find that in order to produce correct pairing, the persistence length of the chromatin fiber, *l*_*p*_, needs to be comparable or greater than the gap size in the button barcode. Given that both *l*_*p*_ and the genomic separation between neighboring ChIP-seq peaks of APs are comparable (;S 1 kb), we surmise that a hierarchical organization of somatic homolog pairing is a structural feature unique to sub-megabase scales (Fig. 6B). It is also anticipated that the distribution of AP binding sites decides and modulates the binding specificity of a chromatin segment of a few kb to its homologous locus, and contributes to shaping 𝒪 (10) − 𝒪 (10^2^) kb sized PIDs.

Taken together, by integrating Hi-C data analyses, 3D structure reconstruction, physics-driven modeling, and simple polymer physics arguments, we characterize the 3D architecture and physical basis of somatic homolog pairing in *Drosophila* embryonic cells. At the genome scale, our Hi-C-based diploid genome models reveal robust end-to-end alignment and overall similarity between the 3D structures of maternal and paternal genome within the same cell (Fig. 2). At the TAD scale, domain boundaries are tightly paired with their homologous loci (Fig. 3). Comparative analyses of two distinct button models further indicate that homolog pairing is orchestrated by specific adhesive buttons, which naturally give rise to overlapping boundaries of *cis* and *thom*-homolog domain boundaries. Finally, the specific button model is expected to impose mechanical coupling between paired homologs. Then, it would be of great interest to study how a symmetry between the pairing-induced *cis* domains is perturbed if an allelic-specific genome editing were made. Such a question is amenable to experimental investigations.

## DATA AND CODE AVAILABILITY

The Hi-C datasets of *Drosophila* used in this study are obtained from the GEO repository with following accession numbers: GSE89112 (Kc_167_ cells), GSE121255 (early embryos), GSE121256 (PnM cells), and GSE171396 (early embryos; Micro-C). The boundaries of TADs in *Drosophila* embryos, larval eye discs, and S2R+ cells are available through GEO accesion number GSE171396, GSE136267, and GSE101317, respectively. The epigenetic states of chromatin segments are acquired from Table S3 of Ref. (59). The coordinates of ChIP-seq peaks of APs in *Drosophila* are downloaded from UCSC Genome Browser server (https://hgdownload.soe.ucsc.edu/gbdb/dm6/reMap/). Codes associated with this work can be accessed upon contacting the corresponding authors.

## AUTHOR CONTRIBUTIONS

L.L. and C.H. designed the research. L.L., Y.J., and Y.T. carried out simulations. L.L., Y.J., Y.T., and C.H. analyzed the data and wrote the article.

## ACKNOWLEDGMENTS

L.L. and C.H. gratefully acknowledge insightful discussions with the late Professor Jie Liang, whose pioneering contributions helped shape early biophysical studies of chromosome organization. This work was partly supported by the Zhejiang Provincial Natural Science Foundation, ZSTU intramural grants (LQ22B040001 and 20062226-Y to L.L.), and a KIAS Individual Grant (CG035003 to C.H.) at the Korea Institute for Advanced Study.

## Supporting Material

### A qualitative theory of bubble chain

#### Insulation of cis contacts

As shown in Fig. 6A, a bubble chain is composed of consecutive bubbles, each consisting of two chromatin arcs from materal (red) and paternal (blue) genome. First, we compare the contact probabilities between points (*i, j*) and (*m, n*) on the same chromosome (blue), which are separated by the same genomic distance of *s*, i.e.,|*i* − *j*| = *s* and |*m* − *n*| = *s*. While *m* and *n* are in the same bubble, *i* and *j*, intervened by an button point *o* (black dot), are in the different bubbles (*i < o < j*). For simplicity, we further assume that *o* point is located at the center between *i* and *j*, so that the associated arc lengths are given as |*i* − *o*| = |*o* − *j*| = *s/*2, and that the intra-bubble statistics is independent of each other. The mean squared distance between (*i, j*) and (*m, n*) can be written as 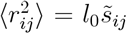 and 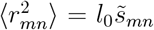 where *l*_0_ is the Kuhn length of the phantom chain, and 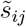 and 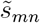 are their effective contour separations. Based on the analogy between an ideal polymer and a random walk^66,67^ and the propagator of a Brownian bridge (see Eq. A2 of Ref. (65)), one finds

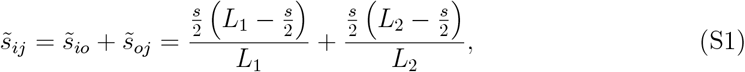

and

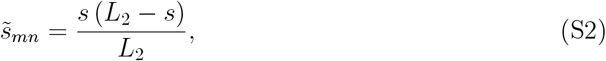

where *L*_1_ and *L*_2_ (see solid gray arrows in Fig. 6A) are the sizes of the two bubbles. It is then straightforward to show

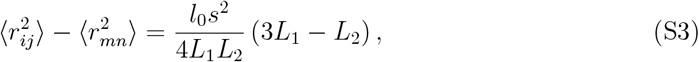

which yields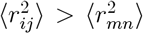. Now, let us consider the case that points (*m, n*) are located in the bubble with a size of *L*_1_, then Eq. S2 becomes 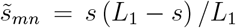.Repeating a similar argument leading to Eq. S3 yields 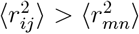. Therefore, as long as the size of two neighboring bubbles are comparable (*L*_2_*/*3 *< L*_1_ *<* 3*L*_2_), the inter-bubble pair (*i, j*) is expected to have a distance greater than that of the intra-bubble pair (*m, n*), which leads to *p*_*ij*_ *< p*_*mn*_ from the anti-correlation between distance and contact probability.

#### Focal contacts at domain corner

Another prominent feature on the map of *cis* contact enrichment of the specific button model is the focal spot at domain corner (Fig. 5B and Fig. S4A). Under a further simplification *L*_1_ = *L*_2_ = *L*, one can get the ratio of the intra-bubble to inter-bubble distance as

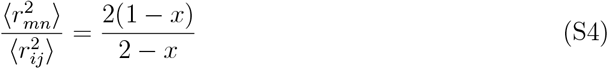

with *x* = *s/L*. Given that the geometrically relevant range of *x* is 0 ≤ *x* ≤ 1*/*2, this ratio decreases monotonically with *x*, which suggests that the maximum enrichment of *cis* contact is acquired at *x* = 1*/*2, i.e., at the corner of the *cis* domain.

#### Modulation of thom contacts near a button

Next, we consider how *thom* contact between a pair of homologous loci (*i, i*^*^) (Fig. 6A) is modulated by the presence of a button point. The mean squared distance between the homologous loci as a function of the genomic distance to the *nearest* button point |*i* − *o*| = |*i*^*^ − *o*| = *d* is calculated as

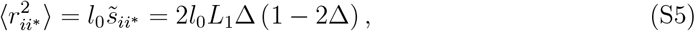

with Δ = *d/L*_1_. As shown in Fig. 6A, the geometrically relevant range of Δ variation is 0 ≤ Δ ≤ 1*/*4, where Δ = 1*/*4 denotes the case when the two loci are at the middle of the arcs comprising the bubble. In this case, the distance 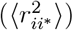 increases monotonically with Δ. In other words, the probability of forming *thom* contact decreases with their arc distance to the nearest button (*d*), and is minimized when the two loci are located at the middle of bubble (Δ = 1*/*4). The button points organize both *cis* and *thom* contacts.

### Supporting Movies

Representative trajectories from Brownian dynamics simulations for (i) the non-specific button model with *l*_*p*_ = 0 (**Movie 1**), *l*_*p*_ = 5.4 (**Movie 2**), *l*_*p*_ = 8*a* (**Movie 3**), and (ii) the specific button model (**Movie 4**). Two pairs of homologous chains (*m*_1_, *p*_1_) and (*m*_2_, *p*_2_) are colored in (red, blue) and (cyan, orange), respectively. The segments colored in gray and black engender the largest gaps and trails shown in Fig. 4B.

**FIG. S1.**
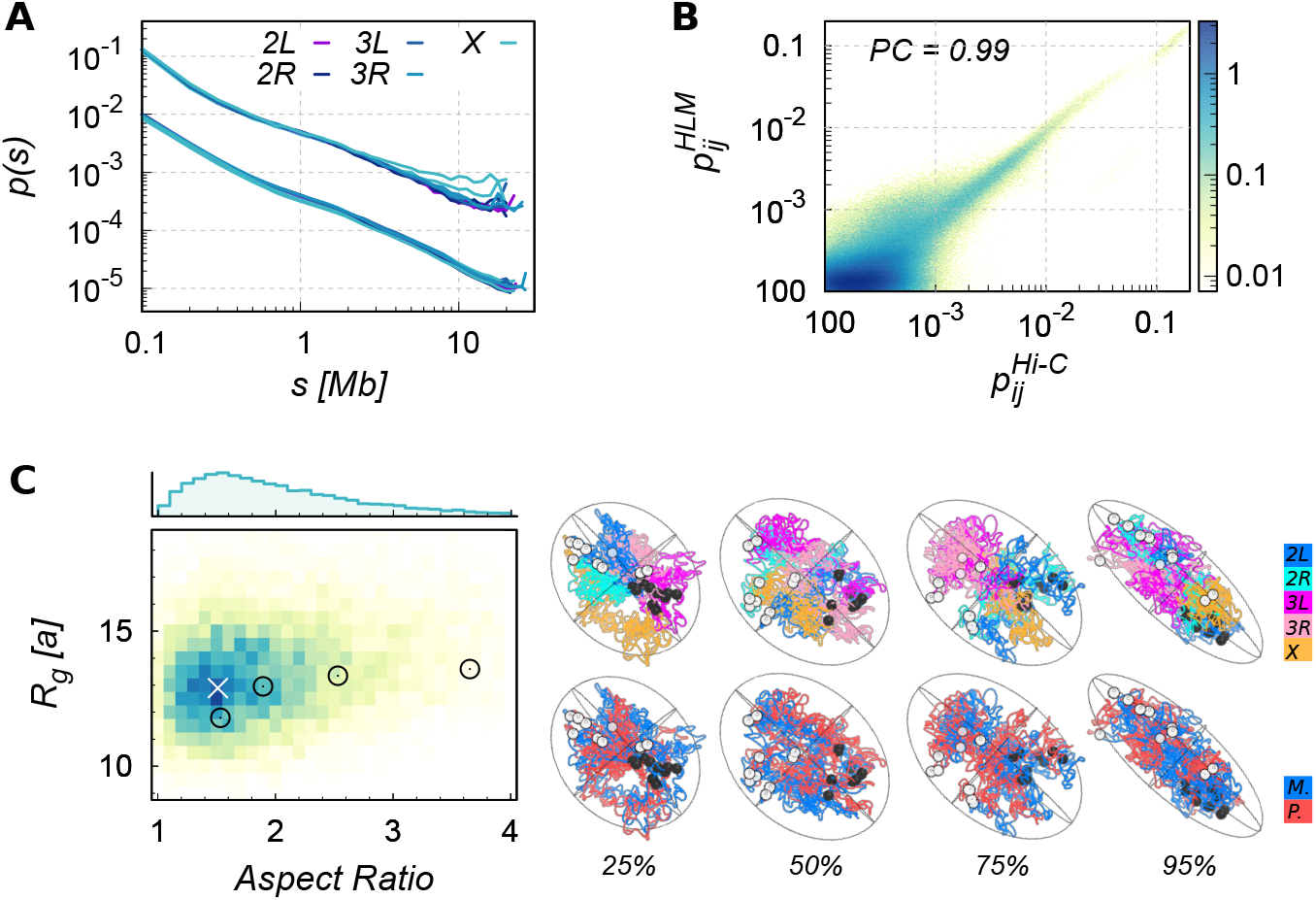
Additional analyses on the HLM of *Drosophila* diploid genome. (**A**) Mean intra-arm contact probabilities as a function of arc length, *p*(*s*). Results from homologous chromosome are plotted with the same colors. Curves based on HLM are vertically shifted (divided by 10) for comparison. (**B**) Correlation between the contact probabilities (*p*_*ij*_) from HLM and from Hi-C, which have a Pearson correlation of 0.99. (**C**) Distributions of the gyration radius (*R*_*g*_) and the aspect ratio of genome in the HLM-based structural ensemble. The most probable structure, at the white cross marker (*×*), is shown in Fig. 2B. Shown on the right are four genome structures at the 25th, 50th, 75th, and 95th percentile values of aspect ratio.

**FIG. S2.**
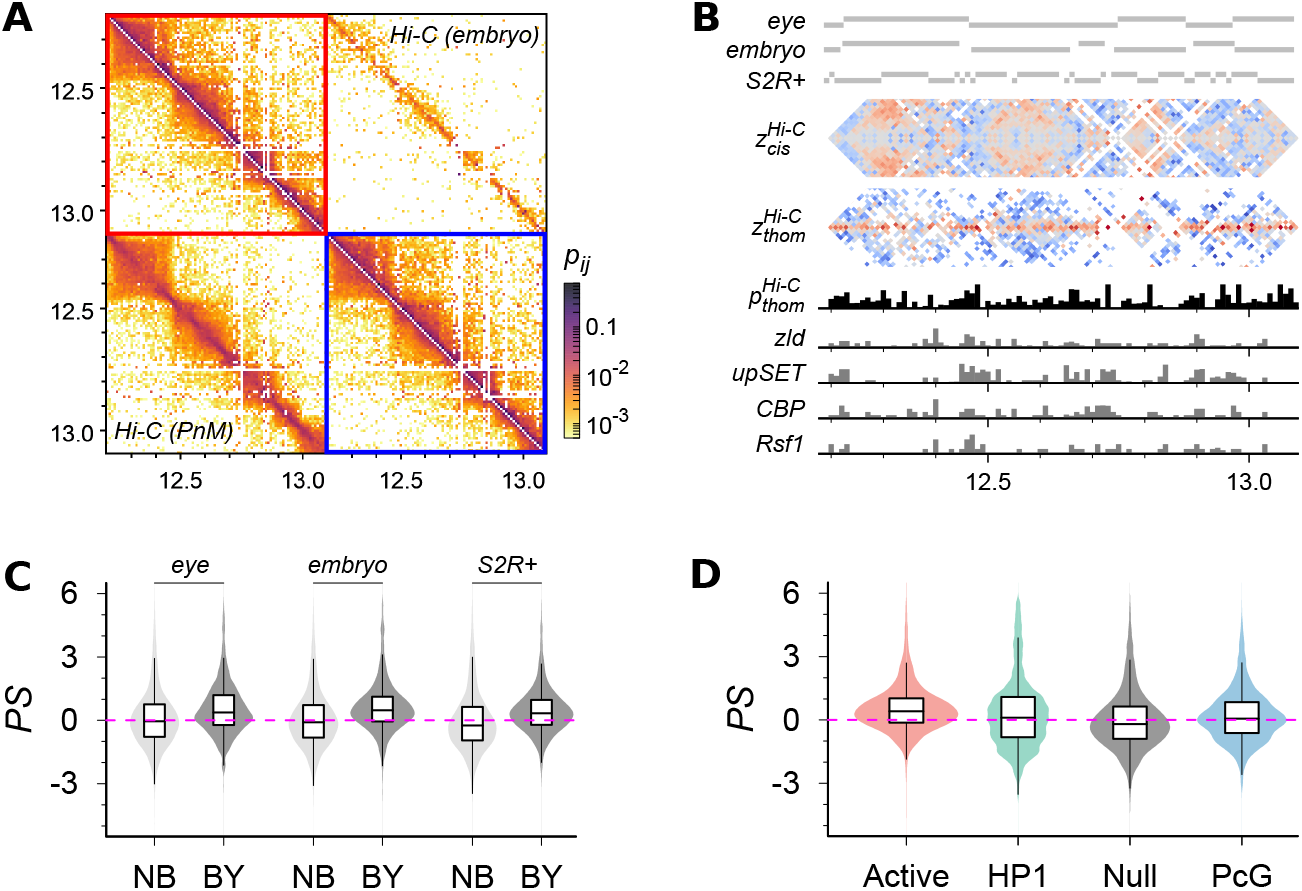
Analyses of *trans*-homolog contacts in early *Drosophila* embryos. (**A**) Haplotype-resolved Hi-C contact map from early *Drosophila* embryos^17^ (upper triangle) versus that from PnM cells^14^ (lower triangle) in a 900 kb genomic region (chr2R:12,200,000-13,100,000) binned at 10 kb. (**B**) Profiles of *cis* TADs called from Micro-C data of *Drosophila* embryos^34^ or Hi-C data of larval eye discs^19^ and S2R+ cells^35^, *cis* and *trans*-homolog contact enrichment 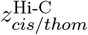 *cis* and *trans*-homolog contact enrichment 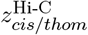 *trans*-homolog contact probability 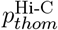 and the number of ChIP-seq peaks of Zelda (zld), upSET, CBD, and Rsf1 proteins^36^ in this region. (**C**) Pairing score, defined as PS = *p*_*thom*_*/* ⟨*p*_*thom*_⟩ where ⟨*p*_*thom*_⟩ is the chromosome-wise mean *trans*-homolog contact probability, of loci at TAD boundaries (BY) and outside the TAD boundaries (NB). (**D**) Pairing score of loci at different epigenetic states^59^.

**FIG. S3.**
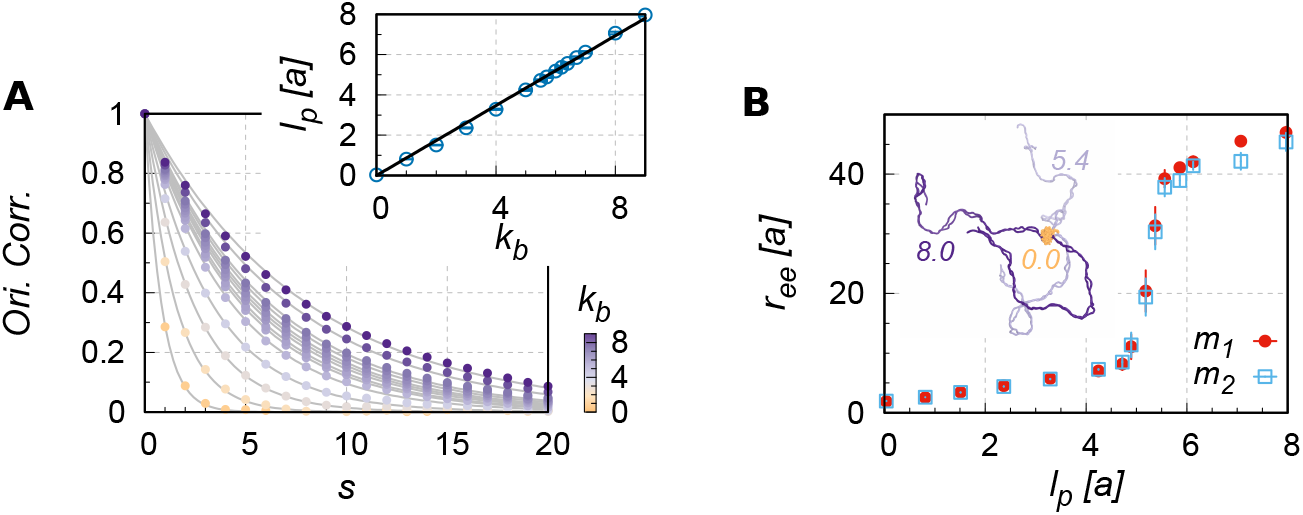
Non-specific button model for two pairs of homologous polymer chains, *m*_1_-*p*_1_ and *m*_2_-*p*_2_. (**A**) Bond orientation (tangent-tangent) correlation 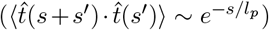 as a function of chain arc length *s* at different values of bending parameter *k*_*b*_. The dependence of *l*_*p*_ on *k*_*b*_ is shown in the inset. (**B**) End-to-end distance of the interacting polymer chains, *r*_*ee*_, versus their single-chain persistence length. Data for chains *p*_1_ and *p*_2_ are omitted, as they are indistinguishable from the *r*_*ee*_ of their homologs *m*_1_ and *m*_2_. Depicted are typical configurations of the system being modeled at *l*_*p*_ = 0, 5.4, and 8.0 *a*.

**FIG. S4.**
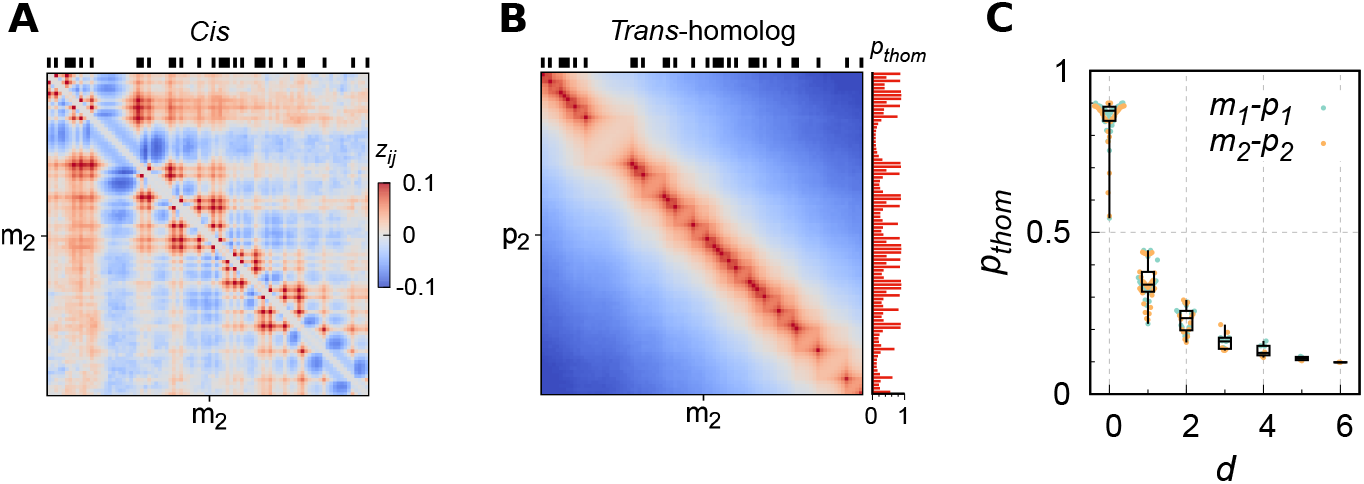
Analyses of the specific button model. Demonstrated are the analysis results on the second pair of homologs, *m*_2_ and *p*_2_. (**A**) *cis* and (**B**) *trans*-homolog contact enrichment maps, with the button barcode on top. (**C**) The *trans*-homolog contact prabability *p*_*thom*_ of monomers versus the arc distance to the nearest button for each monomer (*d*).

**FIG. S5.**
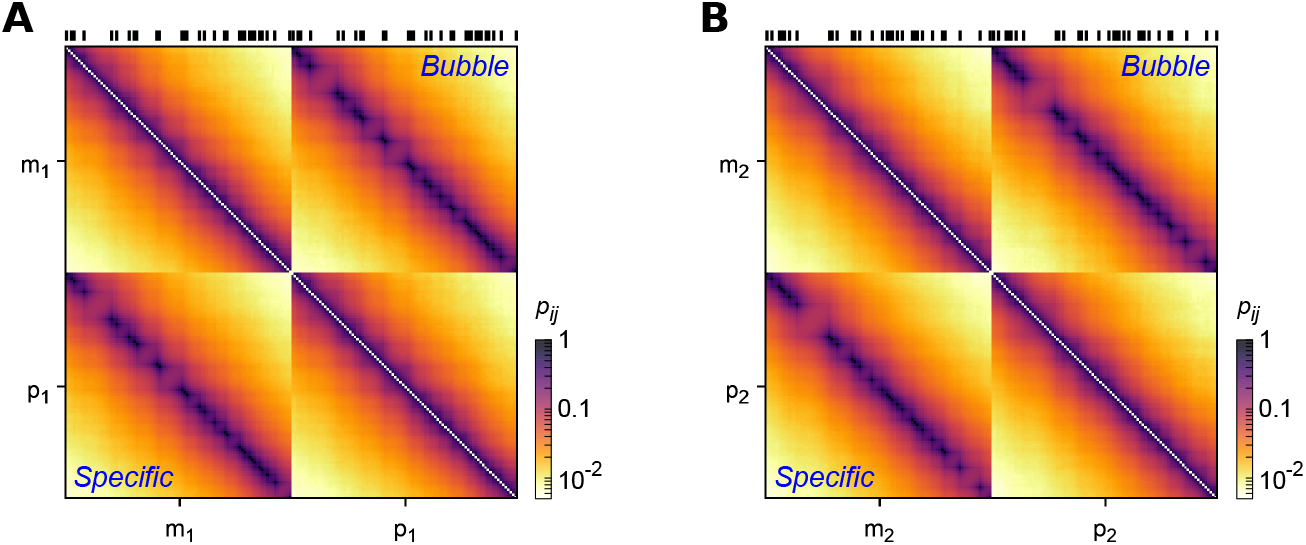
Modeling of bubble chain. As shown in Fig. 6A, instead of a linear polymer, a bubble chain has its chain connectivity (topology) constructed by merging every homologous button pair into a monomer. Brownian dynamics simulations were performed by intergating Eq. 12, with energy potential *U* (***r***) = *U*_bond_(***r***) (Eq. 9). The resulting contact probability maps (upper triagle) for homologs, (**A**) *m*_1_ and *p*_1_ and (**B**) *m*_2_ and *p*_2_, are compared with those from the specific button model (lower triagle). The button barcodes are shown on top.

**TABLE S1.**
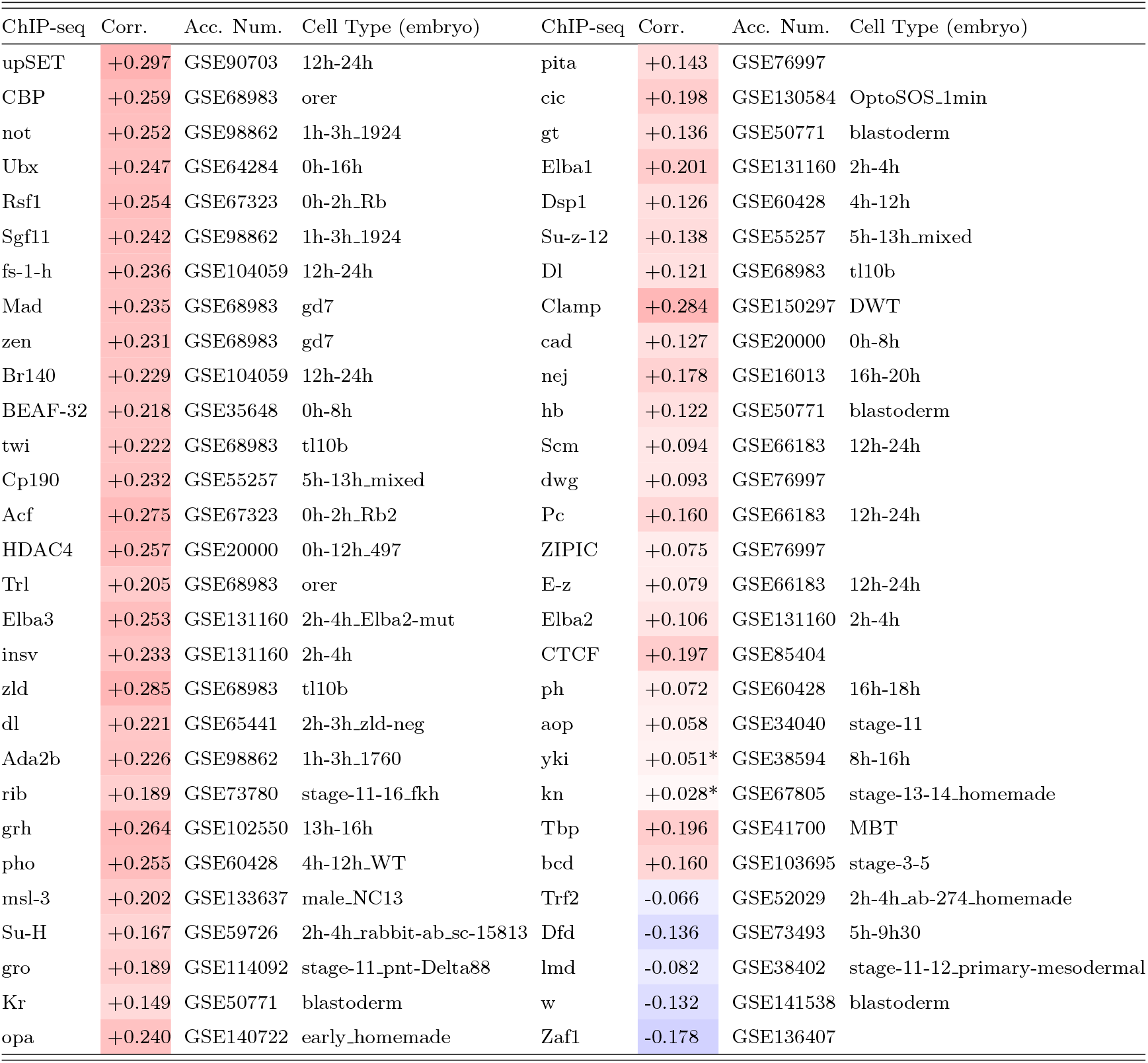
Correlation between protein binding (ChIP-seq peak density) and *trans*-homolog contact probability in *Drosophila* embryos with its value −1 < Corr. < 1 color-coded from blue to red. The names of proteins and cells are the same as in the reMap2022 database^36^. For each protein, the ChIP-seq dataset of the strongest correlation is listed below, while all proteins are ordered by their mean correlation averaged over at least one dataset. The asterisk is appended to the entry with a *p*-value > 1 × 10^−10^. Note that we obtained these results using a bin size of 10 kb. Larger bin size yields overall higher correlations; however, even at 50 kb resolution, the highest correlation is still less than 0.5.

